# Catch and release of sialoglycoRNAs enables sequencing-based profiling across cellular and extracellular material

**DOI:** 10.1101/2025.10.04.680438

**Authors:** Ruiqi Ge, Dennis K. Jeppesen, Sandeep K. Rai, Qin Zhang, James N. Higginbotham, Robert J. Coffey, Ryan A. Flynn

**Affiliations:** Stem Cell and Regenerative Biology Program, Division of Hematology/Oncology, Department of Pediatrics, Boston Children’s Hospital, Boston, MA, USA; Department of Medicine, Vanderbilt University Medical Center, Nashville, TN, USA; Department of Cell and Developmental Biology, Vanderbilt University School of Medicine, Nashville, TN, USA; Department of Stem Cell and Regenerative Biology, Harvard University, Cambridge, MA, USA; Harvard Stem Cell Institute, Harvard University, Cambridge, MA, USA

**Author notes:** These authors contributed equally: Ruiqi Ge, Dennis K. Jeppesen.

## Abstract

Glycosylated RNAs (glycoRNAs) represent a recently discovered class of small RNAs, but their systematic characterization has been limited by reliance on metabolic chemical reporters and high RNA input requirements. Here we present rPAL sequencing (rPAL-seq), a sensitive and selective platform for *de novo* discovery of sialoglycoRNAs. rPAL-seq combines enhanced periodate oxidation of sialic acids with a capture–release workflow and optimized library construction using poly(A) extension coupled with template-switching reverse transcription. The method enabled reproducible profiling from less than 100 ng of input RNA, corresponding to less than 2% of the material required by previous approaches. When applied across 13 human cell lines, rPAL-seq identified lineage-associated glycoRNA patterns alongside a conserved core dominated by uridine-rich snRNAs and snoRNAs, with modification signatures implicating glycosylation on acp³U or related uridine-based modifications. Extending to extracellular vesicles and non-vesicular nanoparticles, rPAL-seq revealed secreted glycoRNA profiles distinct from those of the cellular fraction. rPAL-seq provides a robust, scalable strategy for glycoRNA profiling, opening new avenues for studying this emerging biopolymer.

## Introduction

We initially described a subset of mammalian small RNAs (<200 nucleotides) including tRNA, snRNA, and snoRNA, to exist as a hybrid biopolymer, consisting of the RNA covalently linked to a glycan (glycoRNAs). This work leveraged a metabolic chemical reporter (MCR)-based approach that uses N-Azidoacetylmannosamine-tetraacylated (Ac_4_ManNAz), which uncovered and sequenced the first set of transcripts in HeLa cells and human ES cells^1^. This MCR-based approach has since been widely applied to diverse biological systems^2–7^. However, MCRs are known to suffer from significant issues, including incorporation efficiency and bias, that will need to be overcome to expand our understanding of glycoRNA. An alternative biochemical approach which used ER-targeted engineered ascorbate peroxidase (APEX) proximity labeling offers a means to discover ER-localized RNAs, and demonstrated small RNAs present in the ER^8^, but did not evaluate the glycosylation status of the sequenced RNAs. To mitigate the above issues, we previously developed a chemical labeling strategy for sialic acids termed RNA optimized periodate oxidation and aldehyde labeling (rPAL) that oxidizes the vicinal diols to aldehydes, which can then be modified via a subsequent oxime ligation step^9^. rPAL substantially improved glycoRNA labeling by reducing bias and greatly enhancing the enrichment of glycoRNAs and allowed us to discover that the modified base 3-(3-amino-3-carboxypropyl)uridine (acp^3^U) is necessary for N-linked RNA glycosylation; however, while sufficient for mass spectrometry studies, rPAL suffers from a detectable background labeling of non-glycosylated small RNAs, which would distort interpretations of sequencing data. Other methods for imaging glycoRNAs leverage proximity ligation assay (PLA)^3^ or hybridization chain reaction (HCR)^6^ to monitor specific glycan-RNA pairs in space; however these methods require prior knowledge of the RNA sequence to target.

Discovery of new glycoRNA transcripts requires sequencing platforms. One method employed proximity ligation via a horseradish peroxidase (HRP) enzyme localized to the endoplasmic reticulum (ER) to successfully capture and sequence the surrounding transcripts, which found small RNAs enriched in the ER^8^, but without biochemical evidence of their glycosylation status. Chemoenzymatic approaches have been developed as well. Enzymatic oxidation with galactose oxidase (GAO)^10,11^ enables labeling of multiple galactosyl-containing glycoforms, which can subsequently be analyzed by sequencing. This strategy can capture glycoRNAs without sialic acid (asialoglycoRNAs) but we have previously found that the GAO enzyme can cause RNA degradation (preprint^12^). A very recent preprint^13^ described an α-2,8-sialyltransferase–mediated labeling and capture strategy that circumvents metabolic reporters but still depends on click chemistry, with potential efficiency losses from multi-step labeling. These reported methods commonly use approximately 5–50 μg of input RNA, limiting the contexts in which they are applicable. Thus, while several non–MCR-based approaches exist for glycoRNA labeling, a highly sensitive method that maintains RNA integrity during labeling, ensures glycan-specific release, and is optimized for the analysis of small, structured, noncoding RNAs is lacking.

Here we developed <u>rPAL sequencing</u> (rPAL-seq). By optimizing the rPAL labeling of native sialoglycoRNAs, we further refine the reproducibility and consistency of this non-MCR based technique. We then coupled this to a “catch and release” process by capturing rPAL-labeled glycoRNAs via the sialic acid and selectively cleaving them from neutravidin beads with sialidase under native conditions. Released RNA is transformed into Illumina-compatible libraries with an optimized and multiplexable process assisted by poly(A) tailing, rigorous cDNA synthesis, and template-switching oligo (TSO), enabling readthrough and capture of highly structured small RNAs. We evaluate the sensitivity of rPAL-seq, demonstrating reproducible data using as little as 100 ng of small RNA from cell lines corresponding to less than 2% of prior reports (∼50x more sensitive). We then applied rPAL-seq to 13 commonly used cancer cell lines, characterizing the transcripts that are selectively and commonly expressed. Finally, we examined the sialoglycoRNA transcripts enriched in select secreted extracellular vesicle (EV) and non-vesicular extracellular nanoparticle (NVEP)^14^ fractions from two species, demonstrating the ability to define sialoglycoRNAs in low abundance settings and across species. Together, rPAL-seq provides a simple, flexible, and sensitive method for discovering the suite of sialoglycoRNAs from any RNA sample.

## Results

### Rational design of rPAL-seq

High-throughput*, de novo*, and glycan-specific profiling of glycoRNAs poses several technical challenges. First, glycoRNAs are species of low abundance: previous studies^1,9^ indicate that they represent only a minor fraction of small RNAs (<200 nt), which themselves account for <10% of total RNA. Second, small RNAs are enriched in thermodynamically stable secondary and tertiary structures with a high density of modified bases, both of which impede reverse transcription (RT), enzymatic conversion, and epitope recognition. Third, while lectins can provide glycan specificity, their binding efficiency is substantially lower than that of conventional immunoprecipitation. Chemical/enzymatic conversions (like rPAL) can increase sensitivity, but often at the cost of specificity. Finally, MCR-based profiling is not readily applicable to clinical samples, limiting the study of glycoRNAs across diverse biological and pathological contexts.

We reasoned that these challenges could be addressed through a combination of molecular and computational strategies: (1) addition of oxidation inert polyethylene glycol (PEG) could be used as a crowding agent to enhance rPAL efficiency, as is commonly used for enhancing other enzymatic reactions^15^; (2) use of a thermostable, highly processive engineered Moloney murine leukemia virus (MMLV) reverse transcriptase would be expected to enhance read-through across small RNAs, preserving longer inserts for accurate mapping; (3) sensitivity could be further improved by avoiding T4 RNA ligase bias against structured 3’ and 5’ termini through 3’ end poly(A) extension coupled with template-switching RT^16,17^; (4) incorporation of cordycepin triphosphate (CoTP; or 3′ deoxyadenosine triphosphate, 3’ dATP) could block free 3′ hydroxyl groups and prevent side reactions during periodate oxidation used in rPAL; (5) use of a glycan-specific release such as a sialidase enzyme with a heat-inactivated sialidase control release would provide a critical control against sialic acid–independent enrichment; and (6) complementing these molecular strategies with a computational pipeline incorporating unique molecular identifiers (UMIs) was designed to resolve alignment ambiguities and correct for PCR duplicates, thereby improving quantitative accuracy.

### rPAL-seq enables selective capture and release of sialoglycoRNAs

Based on the above rationale, we established an rPAL-seq workflow to selectively profile sialoglycoRNAs (**Fig. 1a**). Small RNAs were poly(A)-extended, 3′-end blocked with 3′ dATP, rPAL-labeled and neutravidin-enriched, followed by sialic acid-specific release and optimized small RNA sequencing (**Methods**). Because existing commercial kits lacked UMIs and were not compatible with the capture–release workflow, we tested in-house conditions for poly(A) extension and template-switching RT. Conditions were selected to maximize signal-to-noise, defined as desired 50–200 bp inserts relative to adaptor dimers and TSO concatemers^18^ (**Extended Data Fig. 1a**). To assess glycan-specific release, we evaluated rPAL-labeled (with biotin) small RNAs without enrichment. Northern blot detection with streptavidin-IR800 revealed ∼75% reduction of the rPAL-derived biotin signal after *Vibrio cholerae* (VC) sialidase digestion, accompanied by a mobility shift toward higher apparent molecular weight consistent with loss of sialic-acid charge^1^ (**Extended Data Fig. 1b,c**). Digestion efficiency exceeded that of a 1:1 mixture of *Arthrobacter ureafaciens* and *Clostridium perfringens* sialidases, likely reflecting the lectin domain of VC sialidase facilitating recognition of rPAL-modified substrate^19^. To increase sensitivity for limited material, we sought to improve labeling at low input (≤100 ng, ∼10% of bulk). Adding PEG8000 (final concentration of 10.7%, w/v) to the rPAL reaction enhanced signal in low-input samples without adverse effects at bulk scale and was adopted for all subsequent experiments (**Extended Data Fig. 1d,e**). To prevent rPAL-installed biotin from being incorporated into RNA 3′ ends, instead of sialic acids (which could cause RNA capture independent of sialoglycans), we added 3′ dATP to the ATP pool during the poly(A) tailing reaction to block reactive 3′ hydroxyl groups. Concentrations were calibrated to avoid premature termination of poly(A) extension while maximizing signal-to-noise (**Extended Data Fig. 1f,g**).

**Figure 1.**
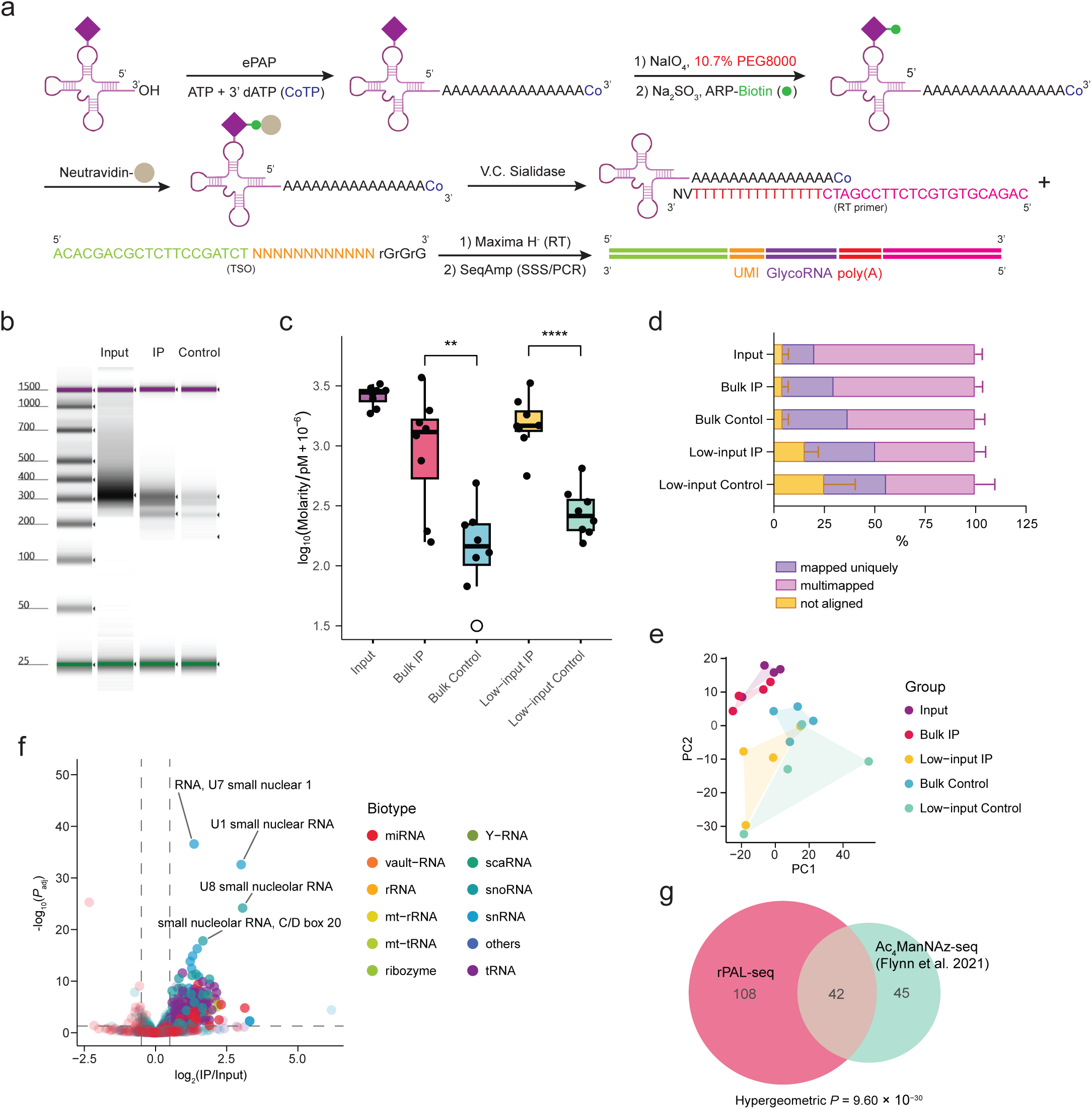
rPAL-seq selectively enriches glycoRNAs and efficiently enables NGS library construction. **a,** Schematic of the rPAL-seq workflow. SialoglycoRNA (represented by a tRNA) was poly(A) extended, 3’ end blocked, rPAL labeled and enriched, followed by sialo-specific release and an optimized small-RNA NGS library preparation workflow. ePAP, *E. coli* poly(A) polymerase; CoTP, cordycepin 5′-triphosphate (3’ dATP); TSO, template-switching oligo; SSS, second-strand synthesis. tRNA icon from BioRender.com. **b,** Representative TapeStation traces of the rPAL-seq libraries. **c,** Quantification of replicates of rPAL-seq TapeStation traces. Concentration was obtained from the area under curve of the region corresponding to 50-200 bp insert. Boxes, interquartile range (IQR); center lines, median; whiskers, 1.5× IQR; points indicate replicates. Out-of-range points are plotted at the y-axis cap (lower, y = 1.5) and shown with an outlined circle. Significance by Student’s *t*-test (*n* = 8, two-sided; \*\**P* < 0.01, \*\*\*\**P* < 0.0001). **d,** Mapping statistics across samples showing fractions of uniquely mapped (purple), multimapped (orange), and unaligned (pink) reads (mean ± s.d., *n* = 8). **e,** PCA of rPAL-seq libraries prepared from HeLa cells. Shaded areas, convex hulls encompassing all replicates of each group. Points indicate biological replicates (*n* = 4 per group). **f,** Volcano plot of glycoRNA families identified by rPAL-seq in HeLa cells. Points are colored by RNA biotype (miRNA, red; vault-RNA, dark orange; rRNA, orange; mt-rRNA, yellow; mt-tRNA, light green; ribozyme, green; Y-RNA, olive; scaRNA, dark green; snoRNA, teal; snRNA, blue; others, indigo; tRNA, purple). Solid points, true positive (TP) hits; transparent points, non-TP. Dashed lines, IP vs. Input thresholds (log_2_FC > 0.5, *P*_adj_ < 0.05). FC, fold change; *P*_adj_, adjusted *P*-value (BH), from DESeq2’s Wald test, two-sided. **g,** Overlap between glycoRNA families detected by rPAL (magenta) and ManNAz (turquoise; Flynn et al. 2021) in HeLa cells. *P-*values by hypergeometric test, two-sided.

Under these conditions, we prepared rPAL-seq libraries from HeLa and OCI-AML3 small RNAs. The libraries showed robust differences between active sialidase release (“IP”) and heat-inactivated mock release (“Control”) across replicates, input scales, and cell types (**Fig. 1b,c, Extended Data Fig. 3a**). Sequencing yielded 70–90% reads mapping to small RNA transcriptome references^20^, with the majority multi-mapped, as expected for short, repetitive, and modified ncRNAs (**Fig. 1d**). A recent benchmark^21^ demonstrated that Expectation–Maximization (EM) based assignment, widely applied in poly(A) RNA sequencing^22,23^ provided the most accurate quantitation of multi-mapping transfer RNAs (tRNAs) compared to commonly used hard-assignment methods. We reasoned that the same principle should extend to other small ncRNAs, and accordingly implemented a custom, UMI-aware, computationally lightweight script (**Extended Data Fig. 2a,** www.github.com/FlynnLab/rPAL-seq). EM-assigned counts clearly separated Input, IP, and Control across scales in HeLa and OCI-AML3 cells (**Fig. 1e, Extended Data Fig. 3c**). Per transcript differential enrichment relative to Input was quantitatively consistent across scales, interestingly not only for IP, but also for Control, underscoring the robustness of the quantification (**Extended Data Fig. 3d,e**).

Next, we used a decision tree to call true positives (TPs) that were not only differentially enriched versus Input, but also significantly enriched above the mock-release control (by Input-normalized contrast of effect size). This strategy excluded both sialidase-independent non-specific captures and cases where IP exceeded Control, but showed only weak enrichment versus Input (“TP_directional”, **Extended Data Fig. 2b**). This yielded various glycoRNA transcripts in HeLa and OCI-AML3 cells (**Fig. 1f, Extended Data Fig. 4; Source Data 1**). The glycoRNA transcript families were well-retained at low-input scale (**Extended Data Fig. 3f,g; Supplementary Table 2**), each contained a range of small nuclear RNAs (snRNAs), small nucleolar RNAs (snoRNAs), tRNAs, microRNAs (miRNAs), short ribosomal RNAs (rRNAs) and Ro60-associated Y5 RNA in HeLa cells which has been individually validated^1^. Systematic comparison with the MCR-based approach showed that rPAL-seq recovered ∼50% of Ac_4_ManNAz-seq hits in HeLa (**Fig. 1g**) and in OCI-AML3 versus H9 (as non-epithelial, **Extended Data Fig. 3h**), a statistically significant overlap that included near-complete recovery of tRNA hits (**Extended Data Fig. 3i,j**). This supports the fidelity of TP identification.

### rPAL-seq reveals recurrent core glycoRNAs and lineage-dependent heterogeneity

We noticed that glycoRNA transcripts identified in HeLa and OCI-AML3 shared many common targets yet appeared distinct on volcano plot: HeLa top hits (by effect size and significance) were enriched for snRNAs/snoRNAs, whereas OCI-AML3 top hits were enriched for tRNAs. This prompted us to perform a systematic profiling of 13 human cell lines, spanning 6 hematopoietic, 5 epithelial, one liposarcoma and one glioblastoma (**Extended Data Fig. 4; Source Data 1, Supplementary Table 2**). Unexpectedly, glycosylation level (expression-weighted, Input-normalized contrast IP versus Control) clustered cell lines into two unsupervised groups broadly matching hematopoietic and non-hematopoietic lineages, and this separation was statistically significant (**Fig. 2a,c**). In contrast, expression level alone produced weaker, non-significant clustering, consistent with the fact that most cell-identity clustering studies have relied on poly(A) RNA expression (**Fig. 2b,d**). We next asked if this lineage-dependent heterogeneity was driven by different repertoires of glycoRNAs, or differential quantitative features driven by a recurrent group of core glycoRNAs. Histograms revealed a long-tailed distribution in either lineage or across all cells combined, showing that most glycoRNAs were cell line–specific, with a subset of core glycoRNAs recurring in ≥75% of cell lines, and a smaller subset universal (**Fig. 2e**). Ranking by recurrence and glycosylation highlighted that the top 20 glycoRNAs were almost exclusively snRNAs and snoRNAs, with tRNAs and rRNAs contributing primarily to hematopoietic lines (**Fig. 2f**). Universally observed glycoRNAs were dominated by U-RNAs, with U1, U2, U4, and U3 ranking among the most prominent hits, indicating that their shared identity extends beyond U-rich sequence composition to near-universal presence of glycosylated copies across cell lines examined (**Fig. 2g; Extended Data Fig. 5a,b**).

**Figure 2.**
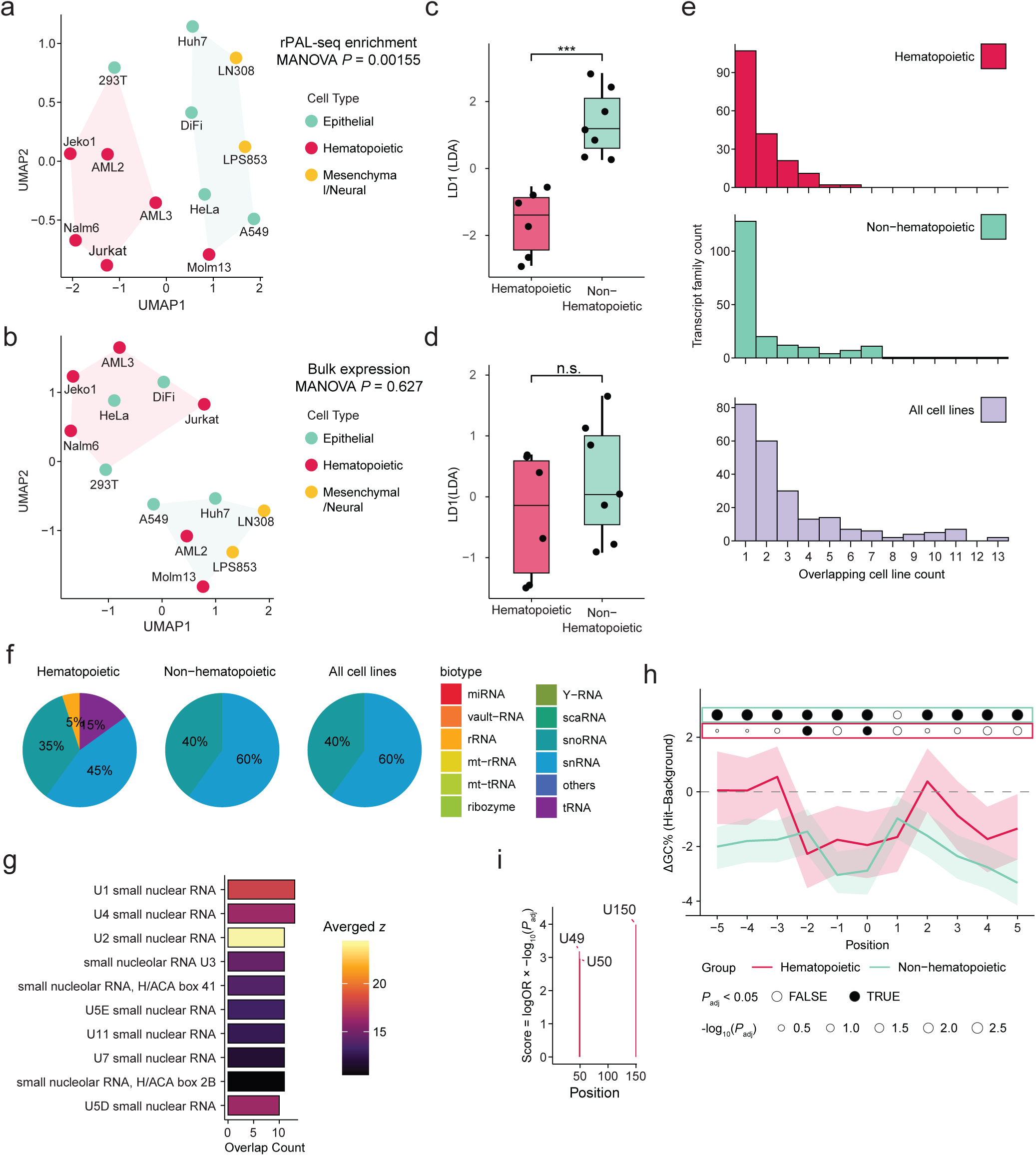
Comparative analysis of glycoRNA signatures across human cell lines. **a,b,** UMAP projection of glycoRNA signatures across a panel of human cell lines (*n* = 13). Embeddings were generated using (**a**) transcript-level *z*-statistics weighted by average log-expression and (**b**) log-expression values alone. Shaded areas, convex hulls of *k*-means clusters (*k* = 2). Point colors indicate *a priori* cell line origin (hematopoietic, magenta; epithelial, turquoise; mesenchymal/neural, orange). Multivariate analysis of variance (MANOVA) was applied to test group separation. **c,d,** Box plots of LD1 scores (separating hematopoietic vs. non-hematopoietic lines) derived by linear discriminant analysis (LDA) from the UMAP embeddings in (**a**) and (**b**), respectively. Boxes, IQR; center lines, median; whiskers, 1.5× IQR; points denote individual cell lines. Significance by Welch’s *t*-test (*n*_1_ = 6, *n*_2_ = 7, two-sided; n.s., not significant; \*\*\**P* < 0.001). **e,** Histogram of glycoRNA transcript family overlap across hematopoietic (top, magenta), non-hematopoietic (middle, turquoise), and all cell lines (bottom, lavender). **f,** Pie charts showing biotype composition of enriched glycoRNA families in hematopoietic, non-hematopoietic, and all cell lines. Color coding consistent with Figure 1f. **g,** Ranked bar plot showing overlap of recurrent glycoRNA families across all cell lines. Bars are colored by a heatmap scale indicating the averaged transcript-level *z*-statistics (Stouffer’s method). **h,** Metagene profile of ΔGC content (vs. biotype and position paired background) aligned relative to the mismatch position (0) for hematopoietic (magenta) and non-hematopoietic (turquoise) cell lines. Shaded areas, 95% confidence intervals. Significance of enrichment indicated by point size (permutation test, two-sided; *P*_adj_, adjusted *P*-value (BH)), with filled circles representing significant (FDR < 0.05). **i,** Positional enrichment profile along small nucleolar RNA U3 in hematopoietic cell lines. Each bar marks a transcript position with significant mismatch enrichment in IP versus Input.

### rPAL-seq implicates uridine-linked modifications via mismatch signatures

Given the prominence of U-RNAs among glycoRNA targets, we considered whether U-based modifications could provide the chemical basis for glycosylation. The only validated linker to date is acp^3^U ^9^, which forms a covalent bond with the glycan moiety and has been described primarily in tRNAs^9,24^. This raised the possibility that acp^3^U may also occur in snRNAs or snoRNAs, or that related U-based modifications capable of glycan attachment remain to be identified. To investigate this possibility, we performed systematic unbiased analysis of mismatch patterns across 13 cell lines, reasoning that the glycosylated base could hinder RT and yield characteristic mismatches in the sequencing alignment. We required either higher mismatch rates in IP than Input or elevated fractions of bulky-modification signatures (skips, gaps, insertions, deletions), as determined by differential testing. These criteria identified IP-enriched mismatch sites in both hematopoietic and non-hematopoietic groups (**Extended Data Fig. 5c; Supplementary Table 4**). Comparing the conversion spectra (observed versus reference base) of these IP-enriched sites revealed a strong enrichment at U reference bases. These sites showed the largest increase in skip/gap/indel events and were the only reference base with enrichment across multiple conversion classes, indicating non-canonical modifications (**Extended Data Fig. 5d,e**). Consistently, aligning sequences to IP-enriched sites (±5 bp, position 0 at the enriched base) revealed local GC depletion compared to a biotype- and position-matched background (**Fig. 2h, Extended Data Fig. 5f,g**). Having established the global enrichment patterns, we next examined specific IP-enriched sites. In U3 snoRNA, enriched sites at U49, U50, and U150 were consistently detected across all cell lines (**Fig. 2i, Extended Data Fig. 5h**). Position 20a of tRNA-Asn-GTT has long been identified to be acp^3^U in both mammalian cells^25^ and *E. coli*^24^. Our analysis successfully recovered this site, further supporting acp^3^U as one of the major glycoRNA linkage bases (**Extended Data Fig. 5i,j**).

### rPAL-seq reveals distinct glycoRNA pattern in extracellular secretion versus cell-surface

Motivated by recent reports of glycoRNA in/on EVs^4,5^, we applied rPAL-seq to extracellular fractions of DiFi (human colorectal) and MDCK (canine kidney) cells, both commonly used in EV studies. Although rPAL-seq is not topologically resolved, we increased granularity by subdividing crude “EV” fractions into highly purified small EVs (sEVs) as well as exomeres and supermeres, two recently discovered amembranous nanoparticles, following a previously established framework^26–28^. Notably, rPAL-seq was able to identify significantly enriched glycoRNAs even with scarce (as low as ∼50 ng of total RNA) input from NVEPs (**Extended Data Fig. 6a; Source Data 2, Supplementary Table 3**). Parallel to our findings in human cell lines, glycosylation level discriminated secreted fractions from the cellular fraction with stronger statistical support than expression level alone (**Fig. 3a–d**). Unexpectedly, overlap between cellular and secreted (sEV, exomere and supermere) glycoRNAs was limited (**Fig. 3e**). In DiFi, cellular glycoRNAs were dominated by snRNAs/snoRNAs, consistent with the pattern observed in other epithelial cell lines, whereas secreted fractions were enriched for tRNAs (**Fig. 3f**). Further subdividing glycoRNAs into cell-unique, shared, or EV/NVEP-unique categories highlighted the distinct composition of the secreted profile, with enrichment of tRNAs, rRNAs, and miRNAs, and depletion of snRNAs (**Fig. 3g,h, Extended Data Fig. 6b**). Notably snRNAs were the top hits in the overall cellular pool and in the shared category yet completely absent from the secreted-unique set. To address the reduced sensitivity from limited material, we performed an alternative directional hit analysis (defined earlier as “TP_directional”, also see **Extended Data Fig. 2b**). This approach identified more putative glycoRNAs (**Extended Data Fig. 6c–f**), but did not alter the central conclusion: secreted glycoRNAs overlapped little with cellular glycoRNAs. The secreted-unique fraction was particularly enriched in tRNAs and miRNAs, and largely depleted in snRNAs (**Fig. 3g,h; Extended Data Fig. 6b**). Northern blot confirmed that secreted glycoRNAs displayed migration patterns similar to their cellular counterparts, consistent with sialic acid–dependent mobility and in contrast to recent MCR-based results^5^ (**Extended Data Fig. 6g**).

**Figure. 3.**
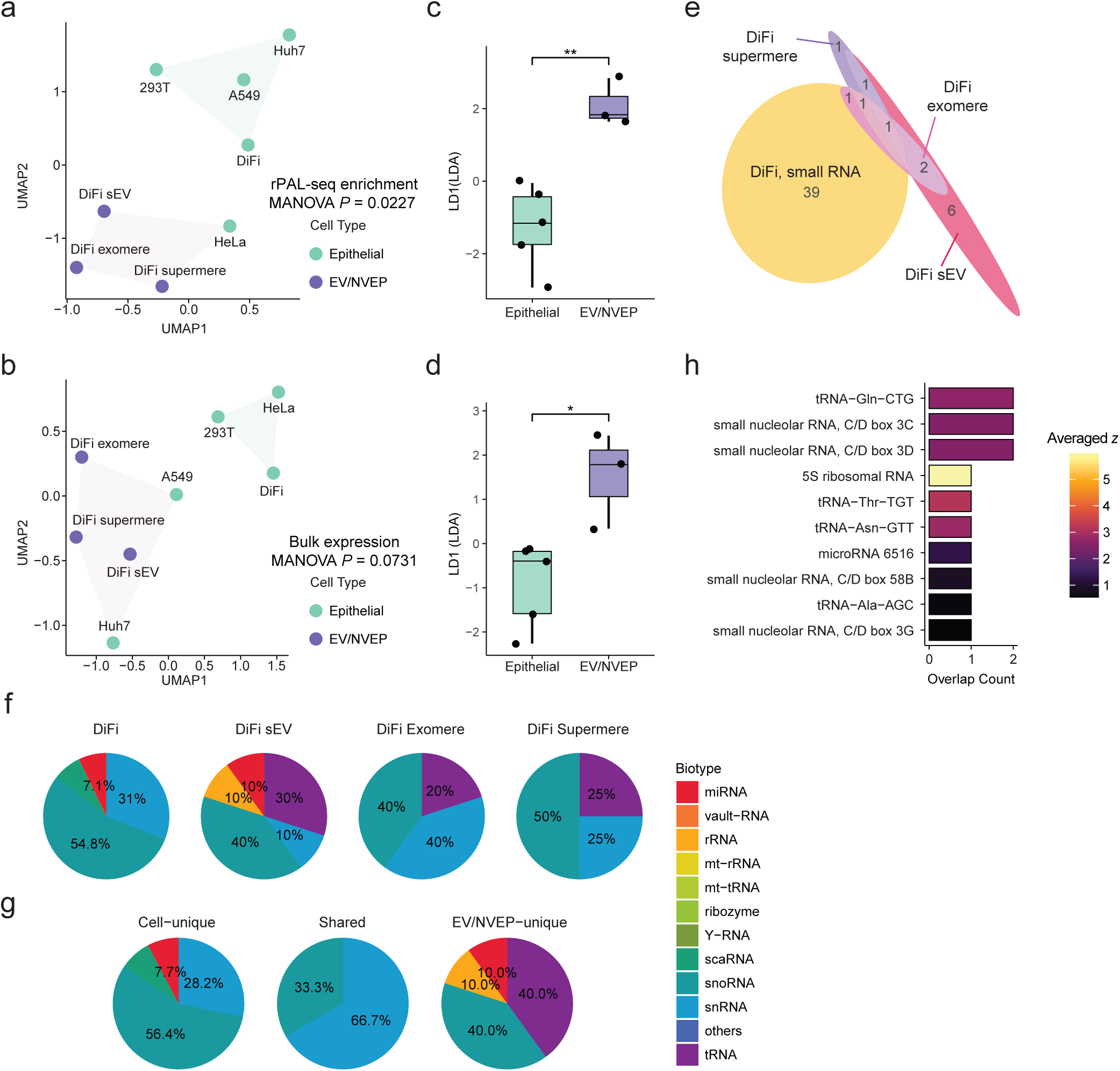
Extracellular glycoRNA profiles are distinct from cellular glycoRNA profiles. **a,b,** UMAP projection of glycoRNA signatures in epithelial cells (*n* = 5) and DiFi cell-derived extracellular fractions (*n* = 3, EV, extracellular vesicle; NVEP; non-vesicular extracellular nanoparticles). Embeddings were generated using (**a**) transcript-level *z*-statistics weighted by average log-expression and (**b**) log-expression values alone. Shaded areas, convex hulls of *k*-means clusters (*k* = 2). Point colors indicate sample category: epithelial, turquoise; EV/NVEP, lavender. MANOVA was applied to test group separation. **c,d,** Box plots of LD1 scores derived by LDA from the UMAP embeddings in (**a**) and (**b**), respectively. Boxes, IQR; center lines, median; whiskers, 1.5× IQR; points denote individual cell lines. Significance by Welch’s *t*-test (*n*_1_ = 5, *n*_2_ = 3, two-sided; \**P* < 0.05; \*\**P* < 0.01). **e,** Euler’s diagram of enriched glycoRNA families detected in DiFi cells (orange), small extracellular vesicles (sEVs, magenta), exomeres (pink) and supermeres (purple). **f,g,** Pie charts showing biotype composition of enriched glycoRNA families in each fraction, and families categorized as cell-unique, shared, and EV/NVEP-unique. Color coding consistent with Figure 1f. **h,** Ranked bar plot showing secreted unique glycoRNA families. Bars are colored by a heatmap scale indicating the averaged transcript-level *z*-statistics.

We next analyzed MDCK cells of canine origin (**Extended Data Fig. 7a,b; Source Data 2, Supplementary Table 3**). The same dichotomy was observed across species: cellular glycoRNAs were enriched in snRNAs and snoRNAs (again with U1 and U3 among the most prominent hits), while secreted-unique fraction was enriched in tRNAs and largely depleted in snRNAs (**Extended Data Fig. 7c-f**). Notably, the top secreted-unique tRNA was tRNA-Gln-CTG in both DiFi and MDCK samples, highlighting a shared feature of secreted glycoRNA profiles. Motivated by this conservation, we profiled a murine epithelial line, EMT6. Despite differences in sample size and origin, the three species exhibited statistically significant overlap (**Extended Data Fig. 7g**). U1–U4 again ranked among the five most conserved glycoRNAs, with seven of the top ten conserved hits belonging to the U-RNA family (**Extended Data Fig. 7h**).

## Discussion

In summary, we developed rPAL-seq, a sensitive and specific strategy for *de novo* profiling of sialoglycoRNAs. By combining enhanced periodate oxidation with a catch–release workflow and optimized library construction based on poly(A) extension and template-switching RT, rPAL-seq achieved reproducible profiling from less than 100 ng of input RNA (**Fig. 1**). Application across 13 human cell lines revealed lineage-associated glycoRNA patterns together with a conserved core dominated by uridine-rich snRNAs and snoRNAs, and mismatch signatures implicating acp³U or related uridine-based modifications (**Fig. 2**). Extending rPAL-seq to secreted fractions demonstrated that glycoRNA profiles in sEVs and NVEPs (exomeres and supermeres) were distinct from those in the cellular fraction (**Fig. 3**). Additionally, cross-species analyses indicated a conserved core set of glycoRNAs, highlighting the broad applicability of rPAL-seq in diverse biological contexts.

Methodologically, the most important advance of rPAL-seq over previous approaches is the catch–release strategy, particularly when coupled with heat-inactivated sialidase mock-release as a control. Earlier methods typically called hits based only on differential enrichment relative to input. However, highly structured and modified small RNAs have an inherent tendency to adhere non-specifically to capture beads, and this baseline enrichment can vary with RNA structure and modification state, generating false positives. Simply raising thresholds on effect size or statistical significance cannot fully resolve this issue, as enrichment may still lack biological relevance if it does not reflect glycan attachment. By contrast, the sequential combination of glycan-specific capture and glycan-specific release ensures high specificity. rPAL has several advantages over other reported glycoRNA capturing methods. When compared to MCR-based approaches such as Ac_4_ManNAz^1^ and Ac_4_GalNAz^5^, it does not require pre-incubation of cells with MCR probes, avoiding potential perturbation artifacts and enabling direct application to primary or clinical samples. As for other chemoenzymatic conversions, compared to galactose oxidase^10,11^ rPAL operates under milder conditions that preserve RNA integrity, thereby improving sensitivity. Compared to the *Campylobacter jejuni* α-2,8 sialytransferase (CstII) labeling coupled with click-chemistry in a recent preprint^13^, rPAL coupled with VC sialidase excels in efficiency, as the sialidase exhibits ∼400× times faster kinetics (CstII^29^ k_cat_/k_M_ 0.15 mM^−1^s^−1^, VC sialidase^30^ k_cat_/k_M_ 65.6 mM^−1^s^−1^). This difference is expected because glycosyltransferases generally display slower kinetics than the corresponding glycosidases, owing to cofactor requirements and typically weaker substrate binding. In addition, copper-free click chemistry often suffers from lower conversion rates due to steric constraints of the cyclooctyne, while copper-catalyzed click chemistry induces risks of RNA degradation. By contrast, rPAL relies solely on small-molecule, non-sterically hindered, RNA-compatible reagents. In terms of release, VC sialidase outperformed commercial neuraminidases, likely because its lectin domain not only facilitates recognition of rPAL-modified, biotin-labeled substrates but it also increases the local concentration of bound glycans, thereby enhancing cleavage efficiency. Taken together, the catch–release design of rPAL-seq provides both high specificity and high sensitivity.

This work also established a streamlined, high-sensitivity protocol for small RNA sequencing and small RNA immunoprecipitation sequencing (RIP-seq). Most existing glycoRNA sequencing methods relied on T4 RNA ligase–based profiling, a widely used approach with many commercial options but best suited for miRNAs rather than the structured and heavily modified tRNAs, snRNAs, and snoRNAs that emerged as major glycoRNA classes in this study. In contrast, poly(A) extension followed by template-switching RT is more effective because (1) poly(A) polymerase is less hindered by 3′ terminal structures and unaffected by 5′ structures and (2) longer RNAs are more likely to form terminal secondary structures. Additionally, the reduced mobility of longer RNAs lowers the frequency of productive intramolecular collisions, causing T4 RNA ligase to strongly favor short inserts and thereby limiting efficient profiling of longer small ncRNA species such as snRNAs, snoRNAs, and Y RNAs^31^. Previous methods also lacked optimized reverse transcriptase choice or reaction temperature to promote read-through, often producing truncated cDNAs and therefore ambiguous mappings. Compared to Oxford Nanopore sequencing^5^, short-read NGS provides greater alignment accuracy, depth, and statistical power for mismatch-based modification analysis. Notably, these features are compatible with small RNAs isolated from antibody-based immunoprecipitation experiments, suggesting it could improve sensitivity and reduce bias in RIP-seq applications as well.

rPAL-seq revealed previously hidden features of glycoRNAs. Earlier functional studies treated glycoRNAs either as a pooled population^2^ or broad subcategories^7^, obscuring fine-scale features across transcripts, cell types, tissues, or species. By contrast, rPAL-seq identified lineage-resolved, topologically resolved, and cross-species fine structures in glycoRNA profiles. Of particular interest, semi-quantitative analysis of glycosylation levels revealed lineage clustering that could not be reproduced by small RNA expression alone, at least not at the scale analyzed (n = 13 cell lines). This work also provided the first profiling of glycoRNAs beyond human and mouse, suggesting broader conservation across species. In addition, the high sensitivity of rPAL-seq allowed profiling of scarce secreted material from amembranous NVEPs, extending prior studies of glycoRNAs in sEVs/exosomes^4,5^. Specifically, tRNAs, and to some degree miRNAs, were prominently featured among the glycoRNAs present in sEVs, exomeres and supermeres. Interestingly the imaging study^4^ also found U1 and U3 to be top biomarkers in sEVs for distinguishing cancer from normal tissue, suggesting potential for direct adaptation of rPAL-seq or rPAL-qPCR in diagnostic applications.

Despite these advantages, rPAL-seq has limitations. Firstly, the current method is restricted to sialoglycoRNAs, leaving asialo-glycoforms unprofiled. We anticipate this could be addressed by applying the optimized protocol to alternative capture–release schemes using lectins and other glycosidases. Secondly, mismatch enrichment analysis cannot rule out the possibility of *in-cis* co-enrichment of non-glycan modifications. For example, in a hypothetical scenario where glycoRNAs were consistently enriched in 1-methyladenosine (m^1^A), it would not be possible to distinguish mismatch signatures arising from m^1^A versus glycan attachment. Future application of rPAL-seq to “writer” knockout models (e.g. DTWD2 for acp^3^U) will help disentangle these effects. Finally, while we analyzed a diverse set of cell lines and species, broader sampling across additional cell types and species could not only refine lineage-level resolution (e.g., distinguishing B-from T-cell lymphoma or epithelial from mesenchymal lineages) but also strengthen observations of evolutionary conservation.

---

**Extended Data Figure 1.**
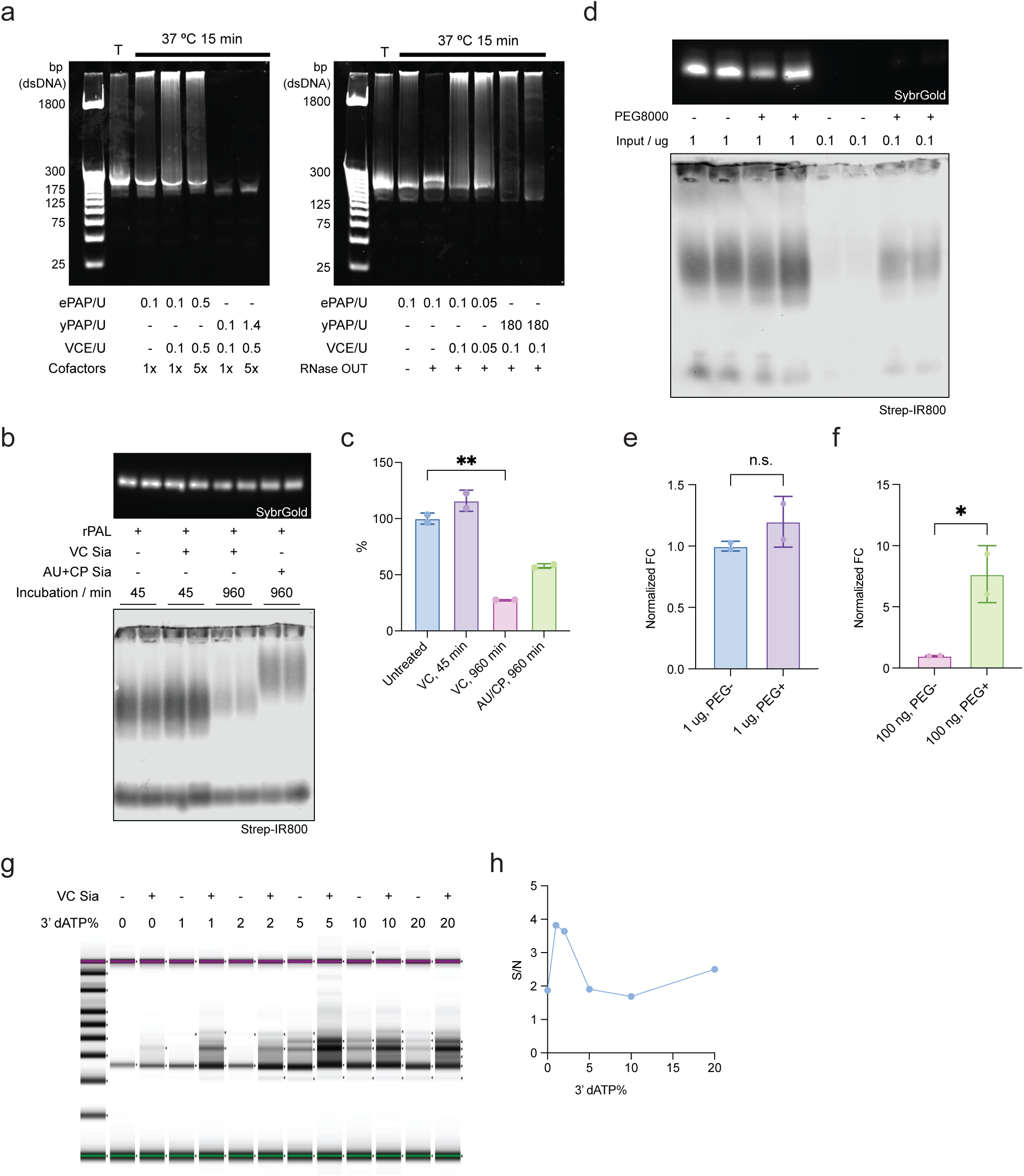
Optimization of rPAL-seq protocol. **a,** Optimization of small RNA NGS library preparation. Libraries are resolved by PAGE and visualized by SYBR Gold. ePAP, *E. coli* poly(A) polymerase; yPAP, yeast poly(A) polymerase; VCE, *Vaccinia virus* capping enzyme. “T”, SMARTer smRNA-Seq Kit for Illumina (Takara). **b,c,** Optimization of sialidase release of captured glycoRNA. (**b**), Top: small RNA resolved by PAGE and visualized by SYBR Gold. Bottom: corresponding Northern blot detection of biotinylated RNA using streptavidin-IR800. (**c**), Quantification of (**b**). VC, *Vibrio cholerae*; AU, *Arthrobacter ureafaciens*; CP, *Clostridium perfringens*. Significance by Student’s *t*-test (*n* = 2, two-sided; \*\**P* < 0.01). **d–f,** Optimization of rPAL labeling of low input samples. (**d**), Top: small RNA resolved by PAGE and visualized by SYBR Gold. Bottom: corresponding Northern blot detection of biotinylated RNA using streptavidin-IR800. (**e,f**), Quantification of (**d**), bulk (blue, purple) and low-input (pink, green), respectively. Significance by Student’s *t*-test (*n* = 2, two-sided; n.s., not significant; \**P* < 0.05). **g,h,** Optimization of RNA 3’ end-block. TapeStation traces (**g**) and corresponding quantification (**h**). S/N, signal-to-noise ratio.

**Extended Data Figure 2.**
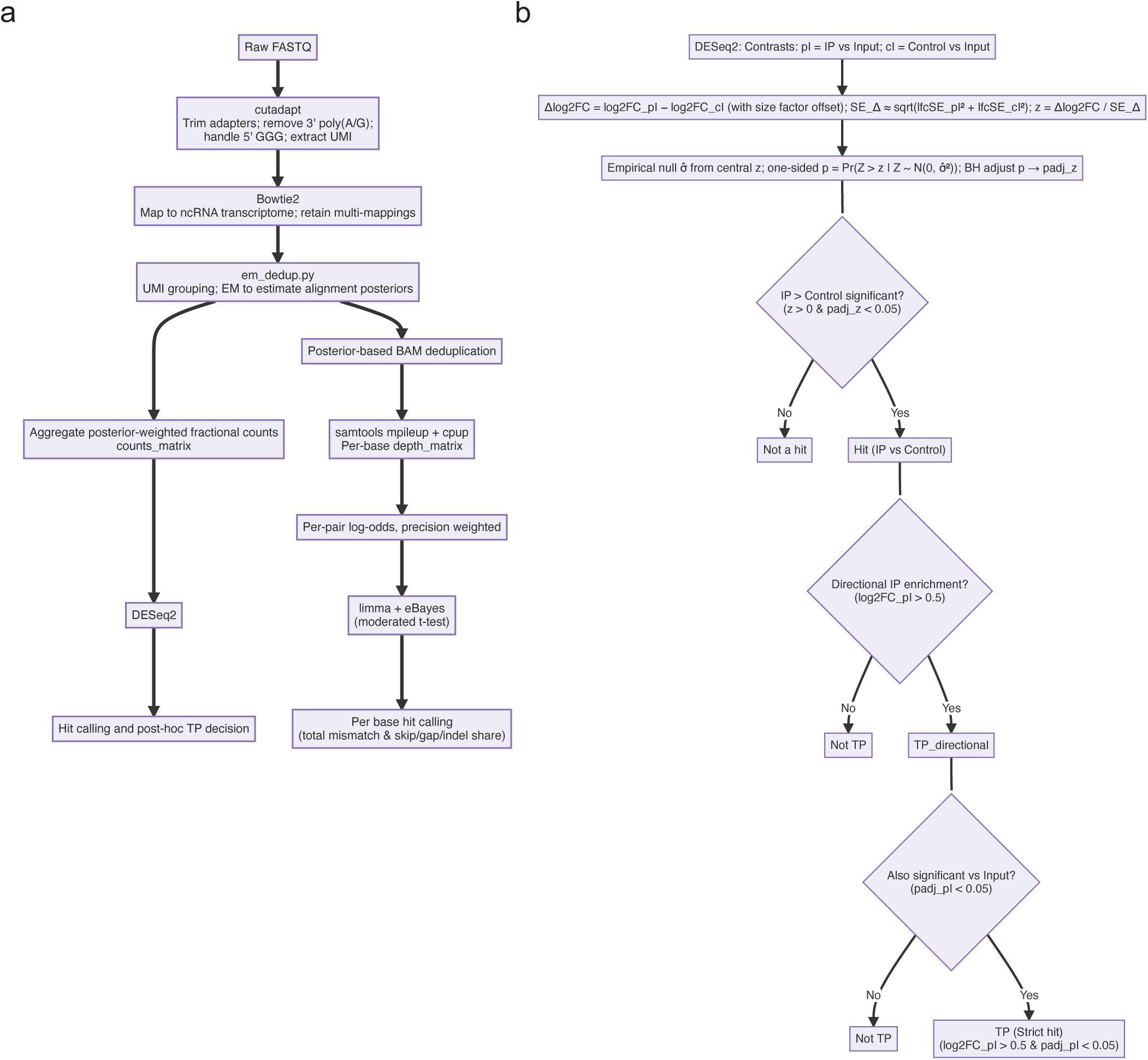
rPAL-seq analysis pipeline. **a,** Computational pipeline of rPAL-seq data analysis. **b,** Decision tree of *post hoc* hit calling.

**Extended Data Figure 3.**
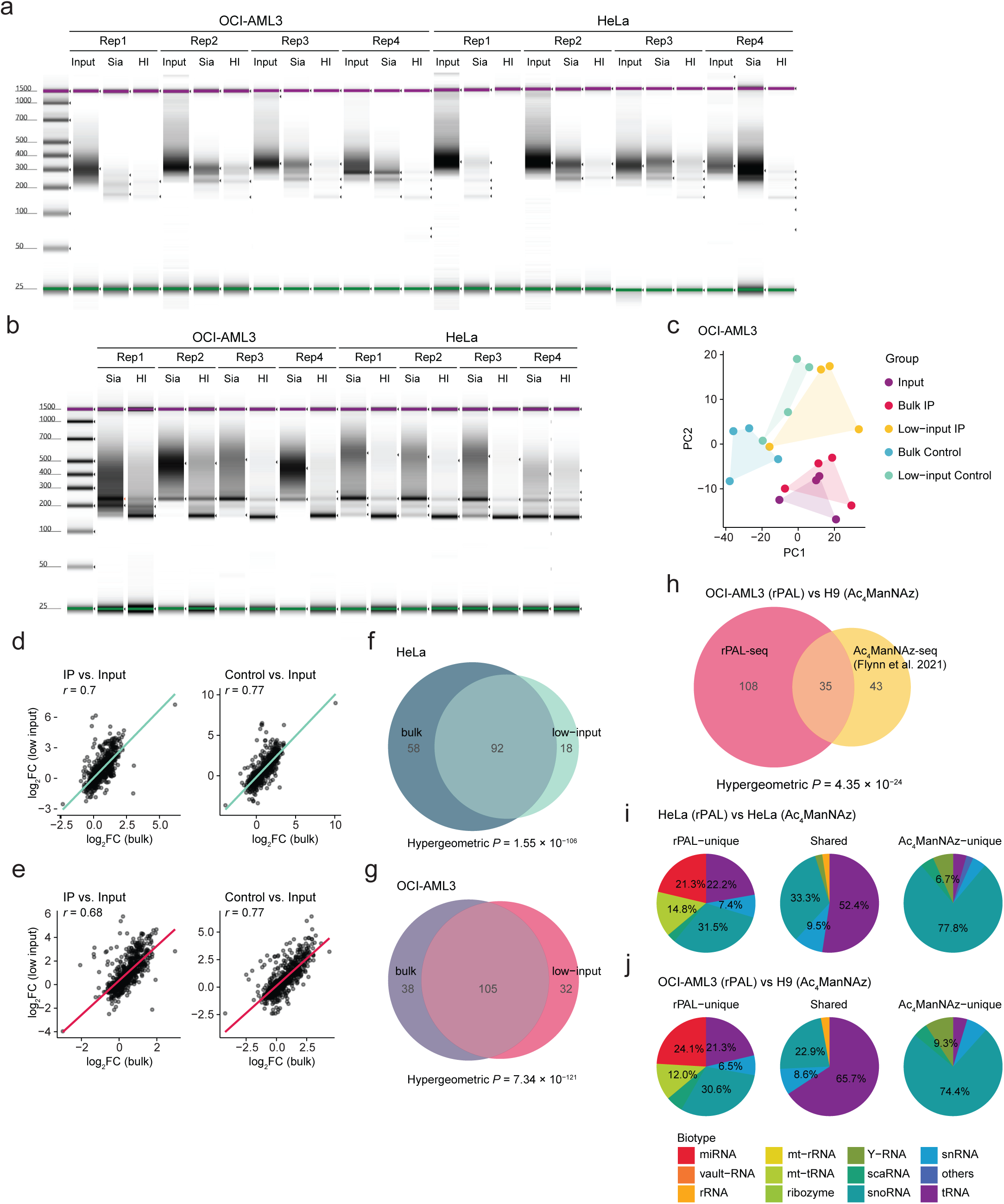
rPAL-seq reproducibility and glycoRNA profiling in OCI-AML3 cells. **a,b,** TapeStation traces of all rPAL-seq libraries prepared from HeLa and OCI-AML3 cells, including bulk (**a**) and low-input (**b**). **c,** PCA of rPAL-seq libraries prepared from OCI-AML3 cells. Shaded areas, convex hulls encompassing all replicates of each group. Points indicate biological replicates (*n* = 4 per group). **d,e,** Scatter plots of bulk vs. low-input comparisons for IP (**e**) and Control (**g**) fractions. *r*, Pearson correlation coefficient, two-sided. **f,g,** Venn diagram showing overlap of glycoRNA families detected in bulk vs. low-input libraries, prepared from HeLa (**f**) and OCI-AML3 (**g**) cells. *P-*values by hypergeometric test, two-sided. **h,** Overlap between glycoRNA families detected by rPAL (magenta) in OCI-AML3 cells and ManNAz (orange; Flynn et al. 2021) in H9 cells. *P-*values by hypergeometric test, two-sided. **i,j,** Pie charts showing biotype composition of enriched glycoRNA families categorized as rPAL-unique, shared, and ManNAz-unique; in HeLa vs. HeLa (**i**) and OCI-AML3 vs H9 comparisons (**j**). Color coding consistent with Figure 1f.

**Extended Data Figure 4.**
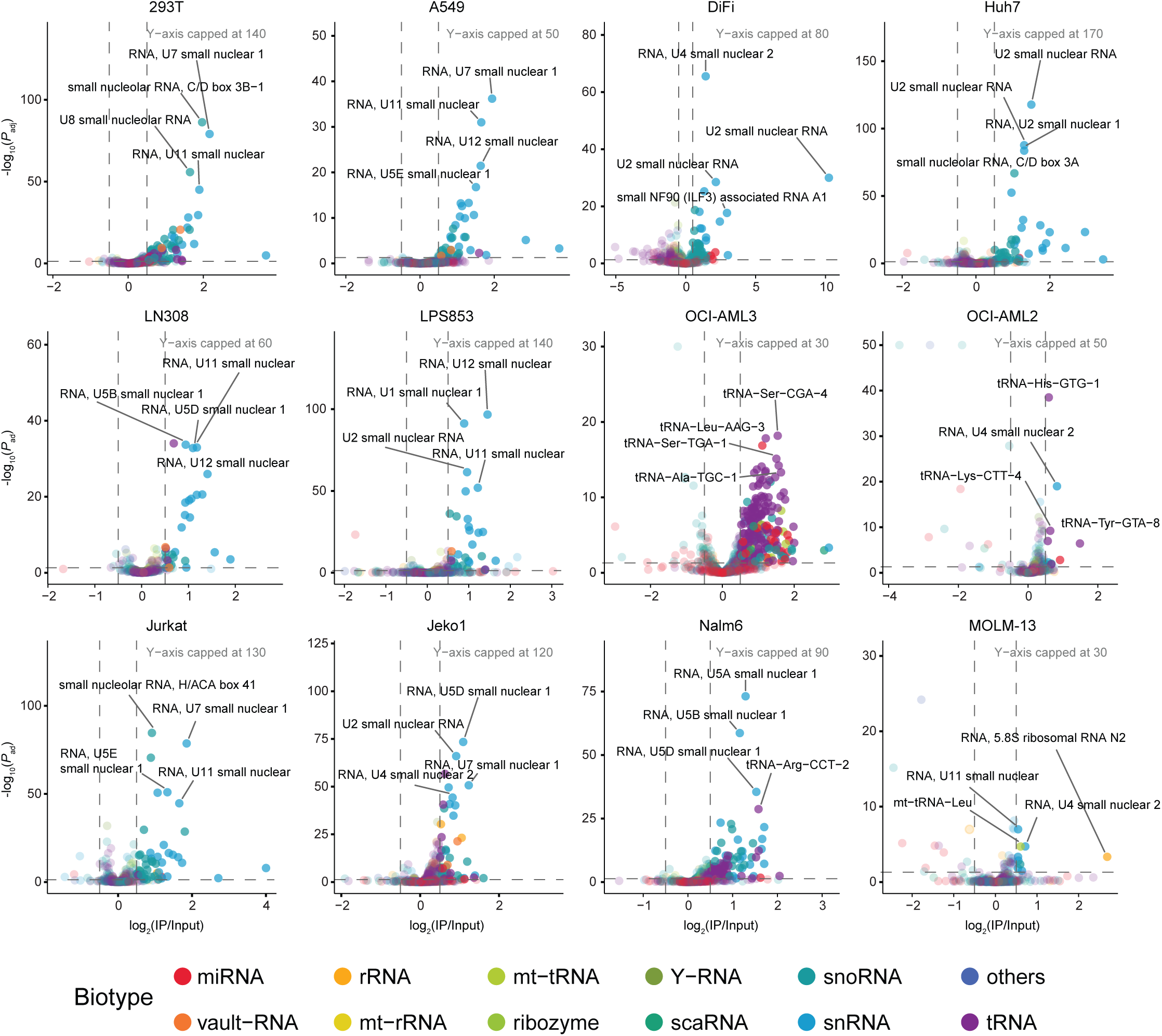
Volcano plots of glycoRNA profiles across 12 cell lines. Color coding consistent with Figure 1f. *P*_adj_, adjusted *P*-value (BH), from DESeq2’s Wald test, two-sided. Out-of-range points are plotted at the y-axis cap and shown with an outlined circle.

**Extended Data Figure 5.**
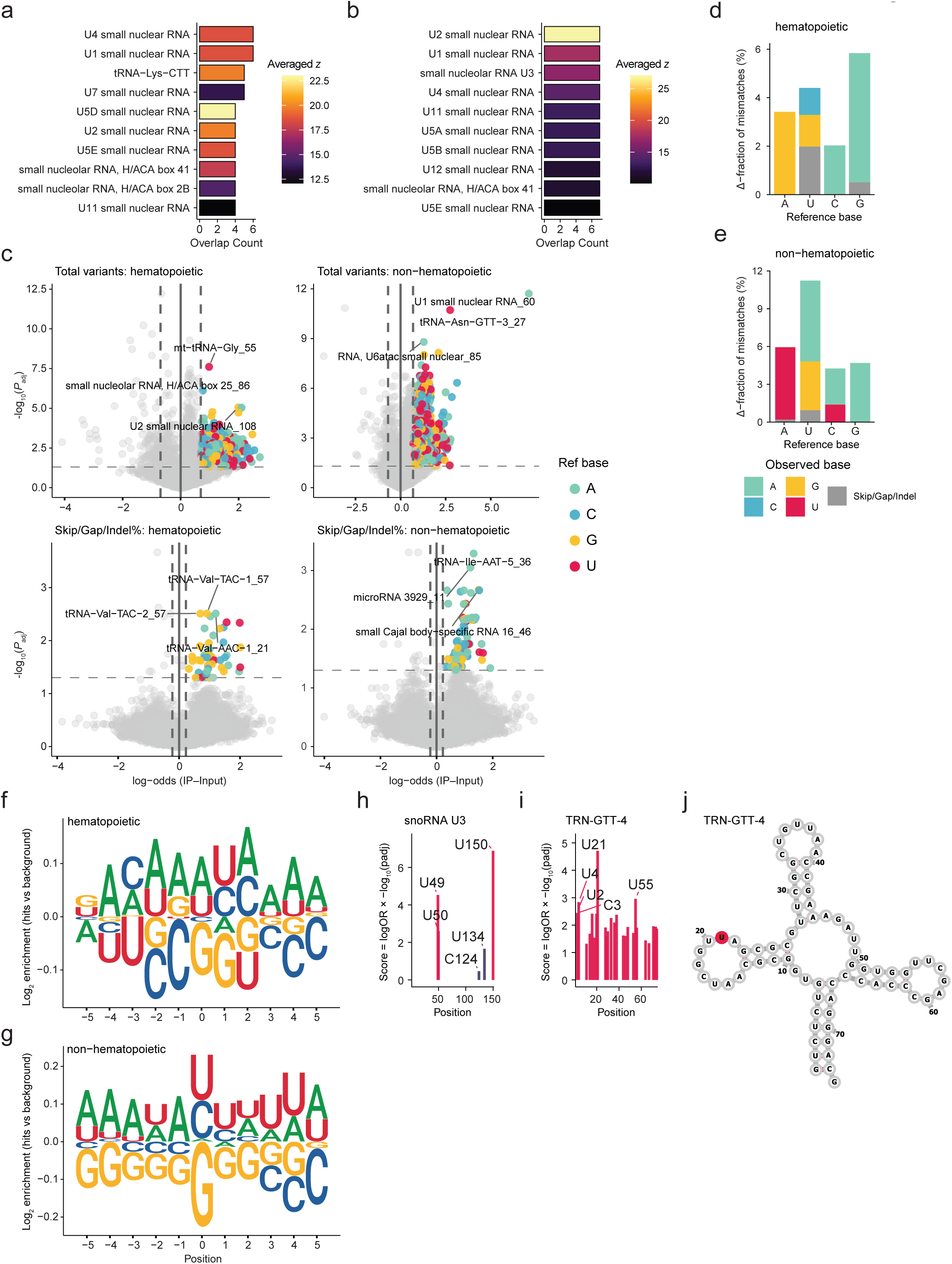
Recurrently enriched glycoRNA families and analysis of IP-enriched mismatches. **a,b,** Ranked bar plot showing overlap of recurrent glycoRNA families across hematopoietic (**a**) and non-hematopoietic (**b**) cell lines. Bars are colored by a heatmap scale indicating the averaged transcript-level *z*-statistics. **c,** Volcano plots of mismatch enrichment in hematopoietic (left) and non-hematopoietic (right) cells. Top, all detected variants; bottom, enrichment in share of skip, gap, insertion/deletion (indel) events. Point colors indicate reference (Ref) base. *P*_adj_, adjusted *P*-value (BH), from limma’s moderated *t*-test, two-sided. **d,e,** Base composition of IP-enriched mismatches, expressed as the Δ-fraction (IP–Input) of observed conversions stratified by reference base in hematopoietic (**d**) and non-hematopoietic (**e**), Colors denote observed base. **f,g,** Metagene sequence logos (vs. biotype and position paired background) aligned relative to the mismatch position (0) for hematopoietic (**f**) and non-hematopoietic (**g**) cell lines. **h,i,** Positional enrichment profile along small nucleolar RNA U3 in non-hematopoietic cell lines (**h**), and profile along tRNA-Asn-GTT-4 in hematopoietic cell lines (**i**). Each bar marks a transcript position with significant mismatch enrichment in IP versus Input. Bar color: magenta, all detected variants; purple, enrichment in share of skip, gap, insertion/deletion (indel) events. **j,** Secondary structure model of tRNA-Asn-GTT-4 showing acp^3^U at position 21 (magenta).

**Extended Data Figure 6.**
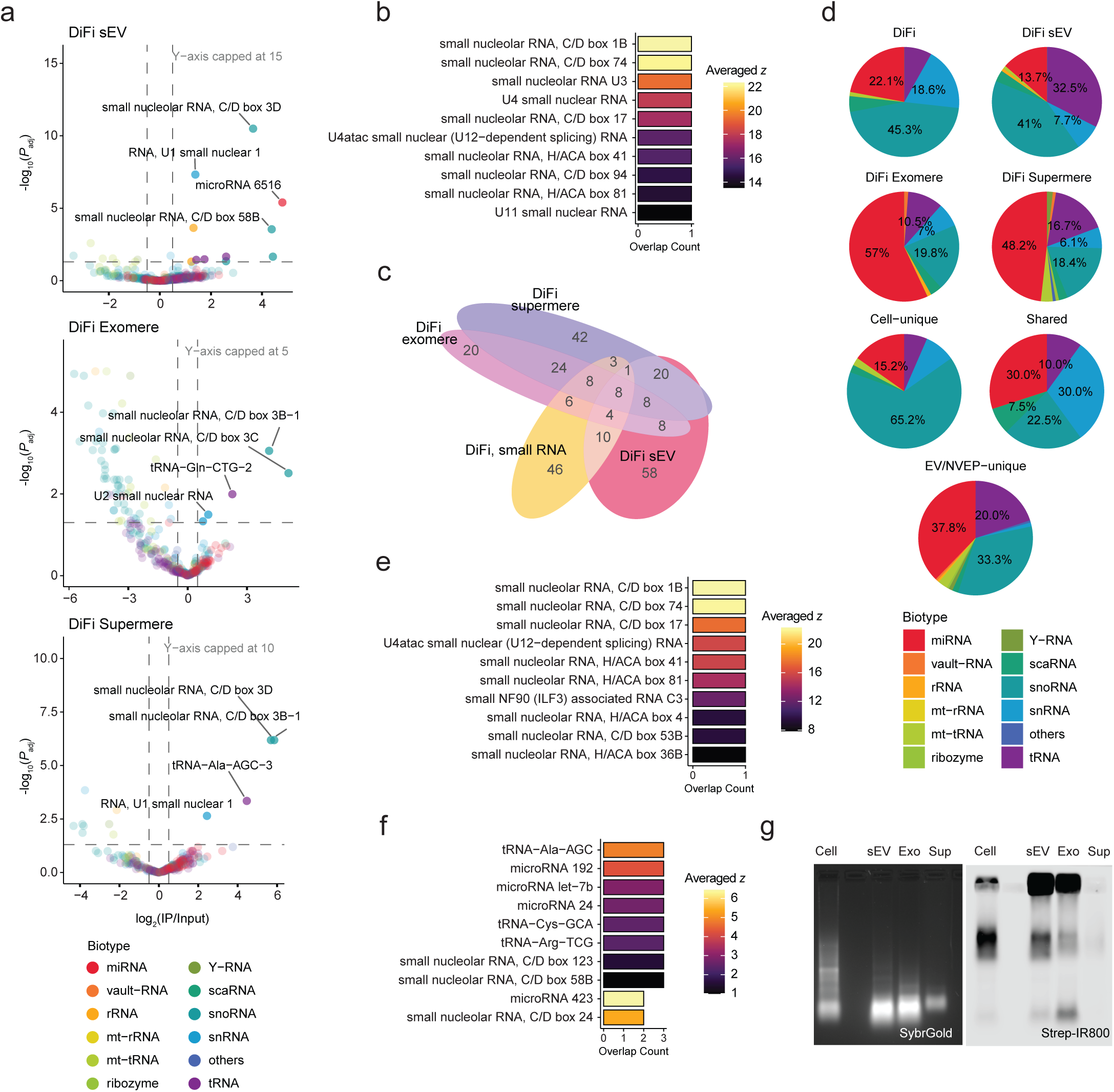
DiFi cell-derived secreted glycoRNA profiles, including alternative directional-hit analysis. **a,** Volcano plots of glycoRNA profiles across DiFi cell-derived secreted fractions. Color coding consistent with Figure 1f. *P*_adj_, adjusted *P*-value (BH), from DESeq2’s Wald test, two-sided. **b,** Ranked bar plot showing cell unique glycoRNA families. Bars are colored by a heatmap scale indicating the averaged transcript-level *z*-statistics. **c,** Alternative analysis using relaxed criteria (“TP_directional” in Extended Data Figure 2b). Euler’s diagram of enriched glycoRNA families (directional) detected in DiFi cells (orange), small extracellular vesicles (sEVs, magenta), exomeres (pink) and supermeres (purple). **d,** Pie charts showing biotype composition of enriched glycoRNA families (directional) in each fraction, and families categorized as cell-unique, shared, and EV/NVEP-unique. Color coding consistent with Figure 1f. **e,f,** Ranked bar plot showing cell unique (**e**) and secreted unique (**f**) glycoRNA families (directional). Bars are colored by a heatmap scale indicating the averaged transcript-level *z*-statistics. **g,** Gel-based detection of DiFi-derived secreted glycoRNA. Left: small RNA resolved by PAGE and visualized by SYBR Gold.Right: corresponding Northern blot detection of biotinylated RNA using streptavidin-IR800. Exo, exomeres; Sup, supermeres.

**Extended Data Figure 7.**
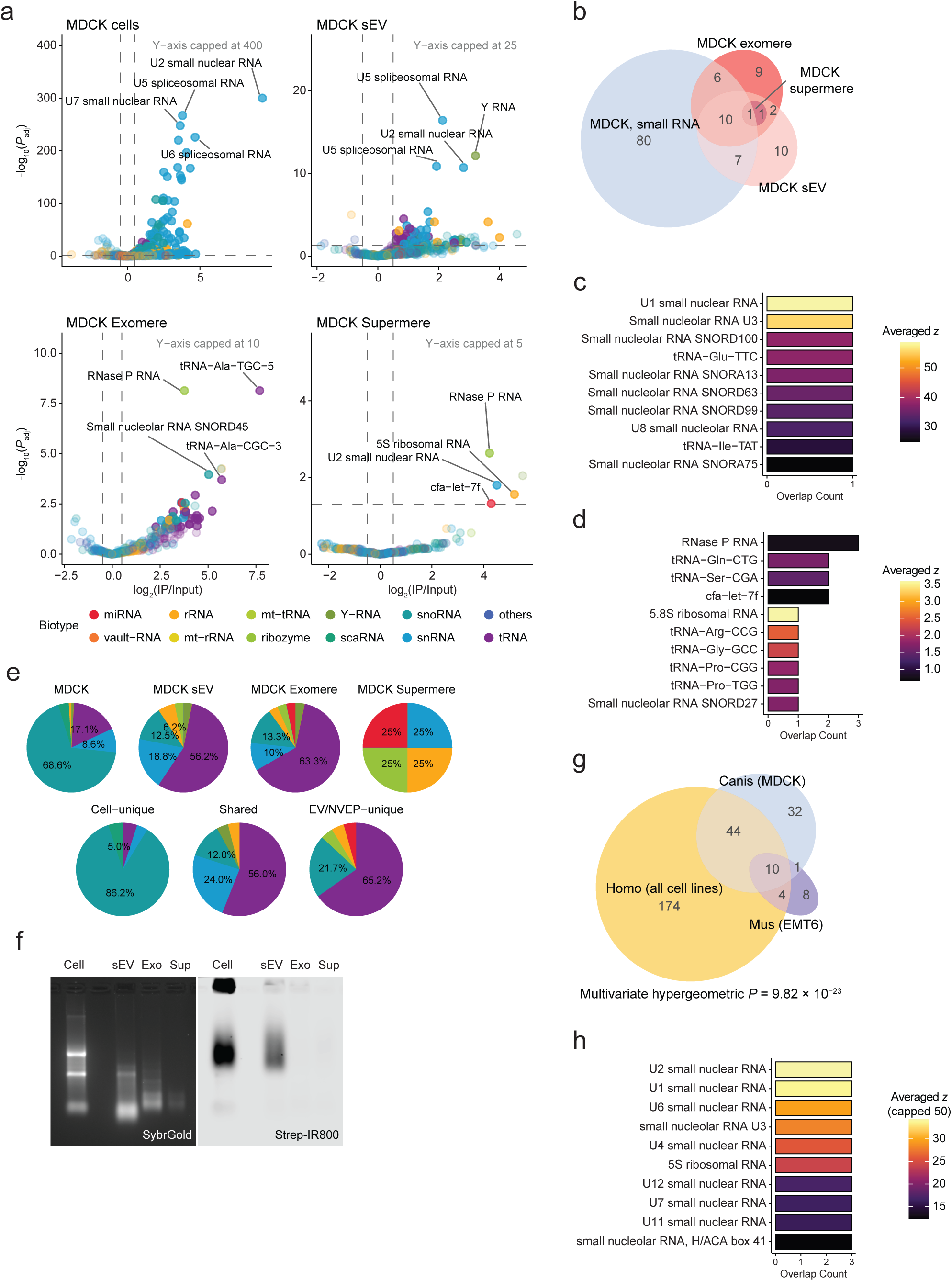
MDCK cell-derived secreted glycoRNA profiles and cross-species comparisons. **a,** Volcano plots of glycoRNA profiles across MDCK cells and MDCK cell-derived secreted fractions. Color coding consistent with Figure 1f. *P*_adj_, adjusted *P*-value (BH), from DESeq2’s Wald test, two-sided. **b,** Euler’s diagram of enriched glycoRNA families detected in MDCK cells (light blue), small extracellular vesicles (sEVs, light pink), exomeres (crimson) and supermeres (dark red). **c,d,** Ranked bar plot showing cell unique (**c**) and secreted unique (**d**) glycoRNA families. Bars are colored by a heatmap scale indicating the averaged transcript-level *z*-statistics. **e,** Pie charts showing biotype composition of enriched glycoRNA families in each fraction, and families categorized as cell-unique, shared, and EV/NVEP-unique. Color coding consistent with Figure 1f. **f,** Gel-based detection of MDCK-derived secreted glycoRNA. Left: small RNA resolved by PAGE and visualized by SYBR Gold.Right: corresponding Northern blot detection of biotinylated RNA using streptavidin-IR800. Exo, exomeres; Sup, supermeres. **g,** Euler’s diagram of enriched glycoRNA families across species: *Homo*, as all cell lines (orange); *Mus*, as EMT6 cells (purple); *Canis*, as MDCK cells (light blue). *P-*values by exact multivariate hypergeometric test, two-sided. **h,** Ranked bar plot showing conserved glycoRNA families across species. Bars are colored by a heatmap scale indicating the averaged transcript-level *z*-statistics (capped at 50).

## Methods

### Cell lines and cell culture

All cells were obtained from American Type Culture Collection (ATCC) unless otherwise specified. The Coffey lab participated in the development of the DiFi cell line; it was authenticated using short-tandem-repeat analysis. OCI-AML2, OCI-AML3, Molm13, Jurkat, Jeko1, Nalm6, and EMT6 cells were cultured in RPMI-1640 medium (Gibco, cat. no. 61870036); HeLa, A549, 293T, Huh7, LPS853 and LN308 were cultured in DMEM medium (Gibco, cat. no. 10566024). DiFi and MDCK cells were cultured in DMEM medium supplemented with 1% L-glutamine (Gibco, cat. no. 25030081) and 1% non-essential amino acids (Gibco, cat. no. 11140050). All media were supplemented with 10% FBS (R&D Systems, cat. no. S11150) and 1% penicillin-streptomycin (Gibco, cat. no. 15140122). Cells were maintained at 37 °C, 5% CO_2_. All cell lines tested negative for mycoplasma contamination (Universal mycoplasma detection kit; ATCC, cat. no. 30-1012K). For production of small extracellular vesicles (sEVs) and non-vesicular extracellular nanoparticles (NVEPs), cells were cultured until 80% confluent. The cells were then washed three times with PBS and cultured in serum-free medium for 48 h.

### EV and NVEP isolation

Both EVs and NVEPs were isolated from cell-conditioned medium as previously described^26^. Detailed protocols are available^27^. Briefly, the cell-conditioned medium was harvested from cells and centrifuged for 15 min at 1,000 × *g* to remove cellular debris and the resulting supernatant was then filtered through a 0.22 µm polyethersulfone filter (Nalgene, cat. no. 568-0020) to reduce microparticle contamination. The filtrate was concentrated using a centrifugal concentrator with a 100,000 molecular-weight cutoff (Millipore, cat. no. UFC9100). The concentrate then was subjected to high-speed centrifugation at 167,000 × *g* for 4 h in a SW32 Ti swinging-bucket rotor (Beckman Coulter) and the resulting sEV pellet was resuspended in PBS containing 25 mM HEPES (pH 7.2) and washed by centrifuging again at 167,000 × *g* for 4 h. To isolate exomeres, the supernatant collected from the 4 h ultracentrifugation was ultracentrifuged at 167,000 × *g* for 16 h. The resulting pellet was resuspended in PBS containing 25 mM HEPES (pH 7.2) and washed by centrifuging again at 167,000 × *g* for 16 h to obtain exomeres. To isolate supermeres, the supernatant from the pelleting of exomeres was subjected to ultracentrifugation at 367,000 × *g* using a Beckman Coulter SW55 Ti rotor (k factor of 48, Beckman Coulter) for 16 h. The resulting supermere pellet was resuspended in PBS containing 25 mM HEPES (pH 7.2).

### High-resolution (12–36%) iodixanol density-gradient fractionation

High-resolution density gradient fraction of crude sEV pellets to obtain pure sEVs was performed as previously described^28^. Detailed protocol is available^27^. Briefly, Iodixanol (OptiPrep) density media (Sigma, cat. no. D1556) were prepared in ice-cold PBS immediately before use to generate discontinuous (12–36%) step gradients. Crude sEV pellets were resuspended in ice-cold PBS and mixed with ice-cold iodixanol in PBS to obtain a final 36% iodixanol solution. The suspension was added to the bottom of a centrifugation tube and carefully overlaid with iodixanol in PBS, in descending order of concentration, yielding the complete gradient. The bottom-loaded 12–36% gradients were subjected to ultracentrifugation at 120,000 × *g* for 15 h at 4 °C using a SW41 Ti swinging-bucket rotor. Twelve fractions of 1 ml were collected from the top of the gradient. Fractions 1 to 6, containing purified sEVs, were pooled. The sEVs were then washed by 10-fold dilution in PBS and subjected to ultracentrifugation at 120,000 × *g* for 4 h at 4 °C.

### Expression and purification of Vibrio cholerae (VC) sialidase

VC sialidase was expressed and purified as previously described^19^ with modifications. The gene block for N-terminal 6×His–VC sialidase (25–781 aa) was purchased from Integrated DNA Technologies (IDT) and cloned into the pET23a vector using infusion assembly protocol. Plasmid sequence was verified by nanopore sequencing. *Escherichia coli* (*E. coli*) BL21(DE3) standard strain was used for the expression at 37 °C, 220 rpm in LB media. Protein expression was induced when the OD_600_ reached 0.6, with the addition of 0.5 mM isopropyl-β-d-thiogalactopyranoside (IPTG). Cells were further grown for 16 h, at 27 °C. Bacterial cell pellets were harvested by centrifugation at 8000 × *g* for 30 minutes at 4 °C. Pellets were washed with 1× PBS and stored at −80 °C until further use.

VC sialidase purification was done by following standard Ni-NTA purification protocol in native buffer conditions. Briefly, cell pellets were thawed and resuspended in 2× PBS, 10 mM imidazole, supplemented with RNase, DNase, and protease inhibitor cocktail tablet. Cells were lysed by sonication (20 min, 10 s ON, 15 s OFF cycles, at 20% amplitude). After sonication, lysates were centrifuged at 12000 × *g* for 30 min at 4 °C. Supernatant was collected and incubated with the pre-equilibrated Ni-NTA resins (QIAGEN, cat. no. 30410) on roller for 2 h, at 4 °C. After incubation, this solution mix was transferred to column. The column was then washed with 50 mL of wash buffer 1 (1× PBS, 10 mM imidazole), 50 mL of wash buffer 2 (1× PBS, 20 mM imidazole). Finally, protein was eluted with 25 mL of elution buffer (1× PBS, 250 mM imidazole). Eluted protein was further concentrated to ∼4 ml and incubated with 2 ml endotoxin-removal resin (in PBS) for 3–4 h at 4 °C on a roller. Endotoxin free protein was recovered by centrifugation (500 × *g*, 1 min) and resin was washed twice with 2 mL of 1× PBS for complete recovery. Protein solution was further polished by size-exclusion chromatography by using Superdex G200 (10/300; Cytiva, cat. no. 28990944) column. Fractions containing protein were pooled together and SDS-PAGE was run to assess the purity level. Protein concentration was estimated using NanoDrop (Thermo) by measuring A_280_ absorbance (ε_280_ = 130,580 M^−1^cm^−1^). Finally, protein was concentrated to 10 μM stock concentration and stored at −20 °C in small aliquots for future uses.

### 3’ Polyadenylation of RNA with end blocking

Total RNA was extracted and fractionated into large and small fractions as previously described^9^. For EV/NVEP, total RNA was used as is. Total or fractionated small RNA (50 ng–2 µg) was heat-denatured at 72 °C for 3 min and immediately cooled on ice. 20 µL of Polyadenylation reaction was assembled with small RNA, 1× poly(A) polymerase reaction buffer (50 mM Tris-HCl, 250 mM NaCl, 10 mM MgCl_2_; pH 8.1), 5U *E. coli* poly(A) polymerase (New England Biolabs (NEB), cat. no. M0276), 40 U RNaseOUT RNase inhibitor (Invitrogen, cat. no. 10777019) and 1 mM ATP (NEB, cat. no. P0756S; diluted in nuclease-free water) spiked with 1% 3′-dATP (Sigma, cat. no. C9137; dissolved in nuclease-free water). Reaction was incubated at 37 °C for 15 min, then immediately cooled on ice. Purification was carried out by mixing the reaction with 3.5 vol. of buffer RLT (Qiagen, cat. no. 79216), 6 vol. of 100% ethanol, and immobilized on Dynabeads MyOne Silane (Invitrogen, cat. no. 37002D). The beads were then washed twice with 80% ethanol and eluted in nuclease-free water.

### Enhanced rPAL labeling, enrichment, and sialo-specific release

50 µL of rPAL reaction was assembled with polyadenylated RNA, 384 µM mPEG_3_-aldehyde (BroadPharm, cat. no. BP-23750), 360 mM MgSO_4_, 288 mM NH_4_OAc (pH 5.0), and 10.7% (w/v) PEG-8000, then incubated at 35 °C for 40 min. On ice, 0.72 mM ARP-biotin (Cayman Chemical, cat. no. 10009350; dissolved in nuclease-free water) and freshly prepared 0.36 mM NaIO_4_ (Sigma, cat. no. S1878; dissolved in nuclease-free water) were added sequentially, mixed, and incubated for 10 min at 25 °C in the dark, followed by quenching with 1.6 mM Na_2_SO_3_. The mixture was incubated for 5 min at 25 °C and then 90 min at 35 °C in the dark. The reaction was purified with Dynabeads MyOne Silane and eluted in nuclease-free water as described in the previous section. 1% (v/v) of the purified RNA was retained as input for further library preparation.

Neutravidin magnetic beads (Cytiva, cat. no. 78152104011150) were washed in biotin wash buffer (10 mM Tris-HCl, 1 mM EDTA, 100 mM NaCl, 0.05% Tween-20; pH 7.5) and pre-equilibrated with glycogen (Thermo, cat. no. R0551). Purified RNA from rPAL was combined with beads in biotin wash buffer and rotated at 4 °C for 30–60 min. Beads were washed sequentially with (per wash 200 µL): ChIRP buffer (2× SSC, 0.5% SDS; mixed immediately before use) twice, biotin wash buffer twice, NT2 buffer (50 mM Tris-HCl pH 7.5, 150 mM NaCl, 1 mM MgCl_2_, 0.05% NP-40) twice, high-salt wash buffer (5× PBS, 0.1% Triton X-100), low-salt wash buffer (1× PBS, 0.01% Triton X-100), and PBS. Finally, beads were cleared of residual liquid and split to generate positive (immunoprecipitated, IP) and Control groups. Beads-captured sialoglycoRNA in the “IP” group was eluted with VC sialidase (0.1 µM) in 1× GlycoBuffer 1 (5 mM CaCl_2_, 50 mM sodium acetate; pH 5.5) supplemented with 20 U SUPERase·In RNase inhibitor (Invitrogen, cat. no. AM2694). Elution proceeded at 37 °C overnight with end-over-end rotation. A mock elution was performed for the “Control” group with same amount of VC sialidase pre-inactivated at 95 °C for 30 min, in exactly the same buffer and incubation conditions as “IP”. If the starting material (prior to the 3′ polyadenylation step) was below 100 ng, 1 ng of Salmon sperm DNA (Invitrogen, cat. no. 15632011) was added as carrier before overnight digestion. Supernatants were purified by Dynabeads MyOne Silane and eluted in nuclease-free water as described in the previous section. Aliquots of enhanced-rPAL labeled sialoglycoRNA and sialidase digested controls were also processed for northern blot analysis, following previously published protocols^9^.

### rPAL-seq library preparation and sequencing

20 µL of reverse transcription (RT) reaction was assembled by combining rPAL-labeled, sialidase-released glycoRNA with 0.5 µM anchored oligo-dT primer (5Bio-s7-dT15(VN), synthesized by IDT; sequence provided in **Supplementary Table 1**), 5% PEG-8000, and 0.5 mM dNTPs (NEB, cat. no. N0447), denatured at 72 °C for 3 min, and immediately cooled on ice. Reverse-transcription mix was then supplemented with 1× RT buffer (50 mM Tris-HCl, 75 mM KCl, 3 mM MgCl_2_, 10 mM DTT; pH 8.3), 1 mM GTP (NEB, cat. no. N0450), 40 U RiboLock RNase inhibitor (Thermo, cat. no. EO0381), 2 µM template-switching oligo (5Bio-s5-TSOr-N12; sequence provided in **Supplementary Table 1**), and 40 U Maxima H Minus RT enzyme (Thermo, cat. no. EP0752). Reaction was incubated at 42 °C for 90 min, followed by 10 cycles of 50 °C for 2 min and 42 °C for 2 min; then 85 °C for 5 min. RT product was amplified with 0.4 µM unique dual-indexed primer pairs (NEB, cat. no. E6440) in 100 µL PCR containing 1× SeqAmp PCR buffer with 2.5 U SeqAmp DNA polymerase (Takara, cat. no. 638509). Reaction was incubated at 98 °C for 1 min followed by 12–16 cycles of 98 °C 10 s, 60 °C 5 s, 68 °C 10 s. Amplicons were purified by mixing the PCR reaction with 3 vol. of buffer RLT and 1.5 vol. of 100% ethanol, immobilized on Dynabeads MyOne Silane, washed twice with 80% ethanol and eluted in nuclease-free water. Libraries were assessed on the 4200 TapeStation system (Agilent) and concentrations were integrated across 215–365 bp (insert 50–200 bp). Libraries with similar profiles were pooled to desired read fractions based on measured concentrations. Pools were size-selected on a 3% (w/v) low-melting agarose (Fisher, cat. no. BP-165) gel. DNA between 200–500 bp was excised under blue light, then extracted and purified by NucleoSpin gel clean-up kit (Takara, cat. no. 740609) according to the manufacturer’s instructions. Where needed, additional double-sided AMPure XP (Beckman, cat. no. A63880) bead-selection (0.5×–0.9×, v/v) was performed to remove >500 bp and <200 bp fragments. Libraries were sequenced on a NovaSeq X (Illumina) with single-end 100 bp reads and dual 8 bp index reads (Read 1: 100 cycles; Index 1: 8 cycles; Index 2: 8 cycles).

### rPAL-seq reads preprocessing and alignment

Raw FASTQ files were generated by Novaseq X following standard Illumina base calling and demultiplexing procedures. Demultiplexed FASTQ files were processed with `cutadapt`^32^ (v 4.9) via a two-pass procedure to extract 5′ UMI and remove RT derived GGG leaders, long poly-A/G tails, residual Illumina adapters, and low-quality bases, discarding sequences shorter than 15 nt. Trimmed reads were aligned to human or mouse small ncRNA transcriptome^20^ (further curated for better biotype annotation) using `bowtie2`^33^ (v 2.5.4), retaining multi-mappings and preserved alignment score (AS) tags for downstream probabilistic assignment. For the *C. familiaris* small ncRNA transcriptome, raw records were downloaded from RNACentral (https://rna central.org/) and filtered to remove duplicate entries, ambiguous/low quality entries and SNP variants differing by an edit distance of 1. All transcriptome files can be found at www.github.com/FlynnLab/rPAL-seq/data.

### Expectation–maximization (EM)-based multi-mapper resolution

Multi-mapped reads were resolved using an EM scheme adapted from `salmon`^22^/`alevin`^23^. Sequencing records were grouped by unique pairs of UMI and query sequence. Each group *g* was associated with a set of candidate transcripts *C*_ℊ_ supported by retained alignments. Weights were assigned to each candidate transcript *t* ∈ *C*_ℊ_ based on alignment score (AS):

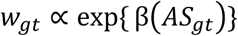

At each iteration (E-step), posterior probabilities were computed as:

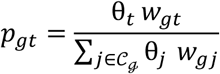

where θ*_t_* is the abundance estimate for transcript *t*. Transcript abundances were then updated (M-step) by:

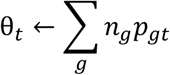

where *n_g_* is the number of sequencing records in group *g*. Iterations continued with light smoothing until the average L1 change fell below tolerance or until the maximum number of rounds was reached. Each group was then hard-assigned to the transcript with maximum posterior probability, with ties resolved by alignment score and lexicographic order.

### SialoglycoRNA differential enrichment analysis

Posterior probabilities from the EM step were used to aggregate posterior-weighted fractional counts per transcript, yielding a transcript-by-sample count matrix. Counts were rounded to the nearest integer, filtered to retain transcripts supported by at least two non-zero observations in both Input and IP samples, annotated with sample metadata (biological origin, condition = Input/IP/Control) and used to construct `DEseq2`^34^ (v 1.42.1) datasets. Size factors were estimated from UMI counts of long rRNAs, under the assumption that these transcripts reflect non-specific binding and remain approximately invariant across samples. Pairwise differential expression (DE) analyses were performed for IP vs. Input and Control vs. Input using `DESeq2`’s Wald test with local dispersion fitting, and results were exported as transcript-level log_2_ fold changes and adjusted *P*-values (Benjamini–Hochberg^35^, BH). DE results from the two contrasts were subsequently merged per group. To assess enrichment beyond the Input control, replicate-matched size factors were used to derive correction offsets. For each transcript, a delta log_2_ fold change and associated *z*-score were computed, with empirical null variance estimated from |*Z*| < 0.5 values. *Post hoc P*-values were obtained from the standard normal distribution and BH adjusted (one-sided). Transcripts were considered true positives (TPs) if they met significance thresholds in both IP vs. Input and IP vs. Control contrasts (FDR controlled at 0.05 each). TPs were further summarized to biologically and sequence-related transcript families. Family-level statistics (log_2_ fold change, adjusted *P*-values) were averaged with weights proportional to transcript baseMean, and combined *z*-scores were obtained using Stouffer’s method. Subsequent general analyses and plotting were performed in R^36^ (v. 4.3.3) using standard statistical and visualization packages, or with Prism 10 (v. 10.4.2).

### UMAP and k-means clustering analysis

For comparative clustering, transcript-level *z*-statistics were weighted by average log-expression (average baseMean from the two `DESeq2` contrasts) and assembled into a group-by-transcript matrix. Transcripts observed in fewer than four groups were excluded. Values were centered and scaled per transcript, and pairwise cosine distances (ignoring missing values) were computed. Classical multidimensional scaling (MDS) was applied to the distance matrix, and the resulting coordinates were embedded with UMAP (min_dist = 0.22, n_neighbors = 10). Groups were clustered in the UMAP space by *k*-means (*k* = 2). Cluster assignments were compared to *a priori* cell type labels (hematopoietic, epithelial, mesenchymal/neural) defined by cell line origin, with convex hulls drawn for visualization. To formally test separation of hematopoietic vs. non-hematopoietic groups, MANOVA was applied to the UMAP1/UMAP2 coordinates, and linear discriminant analysis (LDA) was performed to derive a single discriminant axis (LD1). *t*-tests on LD1 scores were used to quantify statistical significance between classes. The entire procedure was repeated using log-expression values alone under the same parameters to enable direct comparison. Heatmaps of the scaled group matrix were generated using cosine distances for both rows and columns.

### Variant base differential enrichment analysis

After EM-posterior hard-assignment of multi-mapped reads, per-base pileups were generated from BAM files using `samtools`^37^ (v1.21) `mpileup`. Pileup output was processed with `cpup`^20^ (https://github.com/y9c/cpup) to extract per-position base counts. `cpup` outputs were parsed to construct per-position count matrices (reference vs. non-reference bases, insertions, deletions, skips, mismatches), annotated by the same sample metadata used in the previous enrichment analysis section, and extracted for condition pairs IP vs. Input. For each genomic site, mismatches and indels (“total variants”) and total depth were aggregated per condition and replicate. Only sites with ≥20 reads per condition and ≥4 informative replicate pairs were retained. For each pair, log-odds ratios of variant vs. non-variant counts were computed using an empirical logit transform with pseudo-counts, and binomial variances were used as weights. Weighted linear modeling was performed with `limma`^38^ (v.3.58.1), using an intercept-only design, followed by empirical Bayes shrinkage with trend estimation. Per-site statistics (log odds ratio, moderated *t*, *P*-value, adjusted *P*-value (BH)) were reported per supergroup (hematopoietic, non-hematopoietic). In a complementary analysis, the structural component of variants (skips, gaps, insertions, deletions; “SGID”) was evaluated as a share of total variants. Sites with coverage in both Input and IP, and ≥2 informative replicate pairs, were analyzed using the same empirical logit and weighted linear modeling framework. Significant sites were defined using dual criteria: FDR controlled at 0.05 (BH adjusted) and a minimum effect size threshold (log-odds ratio > log 2.0 for total variants, log-odds ratio > log 1.25 for SGID). Filtering was applied separately per supergroup, and hits were exported for downstream analysis. Subsequent general analyses and plotting were performed in R^36^ (v. 4.3.3) using standard statistical and visualization packages.

### Sequence-context analysis of variant sites

For each supergroup (hematopoietic, non-hematopoietic), significant sites (union of “total variants” and SGID) were combined and deduplicated. Background loci were drawn from the same stage files, excluding hits, and matched by genomic position. Centered 11-mers were extracted and per-position nucleotide frequencies computed. The primary analysis quantified per-position ΔGC as the difference between hit GC frequency and a weighted background frequency, with weights reflecting hit counts per position. Uncertainty was estimated by bootstrap resampling of background sequences within strata (B = 500), yielding 95% confidence intervals. Statistical significance was assessed by permutation (B = 500) of hit labels within strata; two-sided *P*-values were calculated per position and corrected across the 11-mer (FDR controlled at 0.05, BH adjusted). Results are shown as ΔGC curves with confidence intervals and significant positions marked. As a complementary analysis, position-specific base enrichments were summarized as log-odds ratios of hit versus background position weight matrices and visualized as sequence logos.

## Data availability

Raw sequencing data (FASTQ files) and processed count matrices generated in this study have been deposited in the Gene Expression Omnibus (GEO) under accession code GSE308686. Source data underlying the figures are provided with this paper (Source Data 1 and 2). Supplementary Tables 1–4 contain identified TP families for all samples and IP-enriched mismatch sites, which also underlie the relevant figures.

## Code availability

All code used for data processing, analysis, and figure generation is available in a GitHub repository (https://github.com/FlynnLab/rPAL-seq; Generative artificial intelligence (ChatGPT-4o and −5; OpenAI) was used to assist script generation and optimization.

## Acknowledgments

We thank Jennifer Porat and others in the Flynn laboratory for helpful comments and discussions. This work was supported by grants from the Burroughs Wellcome Fund Career Award for Medical Scientists (R.A.F.), Rita Allen Foundation (R.A.F.), the Scleroderma Research Foundation (R.A.F.), and the National Institutes of Health, National Institute of General Medical Sciences under award number R35GM151157 (R.A.F.), the National Cancer Institute R35CA197570 and P50CA236733 (R.J.C.). R.J.C. acknowledges the generous support of the Nicholas Tierney Memorial GI Cancer Fund. We acknowledge the Boston Children’s Hospital High-Performance Computing core for supporting the computational analyses of raw sequencing data.

## Author Contributions

R.A.F. conceived of the study. R.G. developed the method, collected the data, performed computational analyses, and generated figures. D.K.J., Q.Z. and J.N.H. generated and purified all the EV and NVEP samples. S.K.R. expressed recombinant VC sialidase. R.J.C. and R.A.F. supervised the study, provided overall guidance, and obtained funding.

